# Multicellular Programs Associated with Right Ventricular Adaptation in Pulmonary Arterial Hypertension

**DOI:** 10.64898/2026.07.30.741916

**Authors:** Catherine E. Simpson, Darin T. Rosen, Andrea Bredemeyer, Haewon Shin, Julie C. Coursen, Sarah L. Khan, Aparna Balasubramanian, Todd M. Kolb, Stephen C. Mathai, Rachel L. Damico, Kathryn C. Fitzgerald, Monica Mukherjee, Kory M. Lavine, David A. Kass, Steven Hsu, Paul M. Hassoun

## Abstract

**Background:** Right ventricular (RV) adaptation determines outcomes in pulmonary arterial hypertension (PAH), yet multicellular molecular programs associated with adaptive versus maladaptive RV remodeling in living humans remain incompletely defined.

**Methods:** We collected 32 human RV tissue biopsies from patients with idiopathic PAH, systemic sclerosis-associated PAH (SSc-PAH), systemic sclerosis without pulmonary hypertension, with 24 nonfailing donor RVs serving as controls. We performed single-nucleus RNA sequencing and integrated cell-type specific transcriptional programs with contemporaneously obtained multi-beat pressure-volume loop measurements of RV contractility (Ees, end-systolic elastance) and RV-pulmonary arterial coupling (the ratio of Ees to Ea, the effective arterial load). SSc modification of PAH-associated biology was assessed using interaction terms. Bulk RV proteomic pathway enrichment was performed to assess an orthogonal molecular layer, and exploratory cell-cell communication analyses alongside independent spatial transcriptomic analyses were performed to contextualize key findings.

**Results:** PAH was associated with broad depletion of biosynthetic, trafficking, and mitochondrial programs across cell types. SSc modified the magnitude of many PAH-associated transcriptional programs while largely preserving pathway directionality. Significant multicellular pathway enrichments were associated with RV-pulmonary arterial coupling. Joint analysis of Ees, Ea, and Ees/Ea identified biologic programs associated with different RV responses to varying loading conditions. Preserved coupling was characterized by enriched extracellular matrix, laminin-integrin, receptor tyrosine kinase, mitochondrial, and translational programs involving fibroblast, endothelial, endocardial, and cardiomyocyte compartments. Cell-cell communication analyses predicted coordinated stromal-vascular signaling networks involving laminin-integrin and endothelial-to-mural signaling in preserved coupling. Proteomic and spatial analyses supported recurrent multicellular themes.

**Conclusions:** RV adaptation in PAH is associated with distinct, coordinated multicellular programs that vary with load and contractile response. RV-PA coupling in PAH is associated with multicellular remodeling that extends beyond cardiomyocytes and reflects organized vascular-stromal support architecture. These findings identify extracellular matrix, laminin-integrin signaling, mitochondrial, and translational programs as associated with adaptive RV remodeling in PAH.

**Clinical Perspective:** *What is new?:*

- Cell type-resolved molecular profiling of living human RV tissue identifies PAH-associated depletion of biosynthetic, trafficking, mitochondrial, and repair-associated programs across multiple cardiac cell types.
- Integration with contemporaneously obtained pressure-volume loop physiology demonstrates that preserved RV-pulmonary arterial coupling under lower load was associated predominantly with cardiomyocyte mitochondrial and metabolic competency, whereas preserved coupling under higher load was associated with extracellular matrix remodeling and vascular-stromal signaling.
- Proteomic, cell-cell communication, and spatial analyses provided orthogonal support for coordinated extracellular matrix and vascular-stromal programs associated with preserved RV-pulmonary arterial coupling.

*What are the clinical implications?:*

- These findings shift the biology of RV adaptation from a predominantly cardiomyocyte-centered model toward a multicellular tissue model in which metabolic, matrix, and vascular support programs vary according to loading conditions and contractile states.
- Extracellular matrix-integrin signaling, endothelial-mural communication, and mitochondrial competency represent candidate pathways for mechanistic investigation toward RV-directed therapies in PAH.

## Introduction

Pulmonary arterial hypertension (PAH) is a progressive pulmonary vasculopathy that presents an increased load to the right ventricle (RV), and consequent RV dysfunction remains the major determinant of symptoms, exercise limitation, and mortality.^1–3^ Current therapies for PAH primarily target the pulmonary vasculature, however outcomes are strongly influenced by the ability of the RV to adapt to increases in load, thereby preserving RV-pulmonary arterial (RV-PA) coupling.^4,5^ Importantly, RV dysfunction in PAH cannot be explained solely by hemodynamic load, as patients with similar pulmonary vascular resistances exhibit markedly different degrees of RV adaptation and thus experience variable clinical trajectories.^6–9^ This heterogeneity is particularly evident in systemic sclerosis-associated PAH (SSc-PAH), in which patients demonstrate disproportionately impaired RV function and worse survival compared to other forms of PAH.^10–13^

These observations suggest that RV remodeling reflects active, coordinated biologic processes within the RV itself. However, the molecular mechanisms underlying adaptive versus maladaptive RV remodeling in PAH remain incompletely defined, particularly across cardiac cell types, as most previous cell-based studies in humans have investigated the mechanical properties of cardiomyocytes.^11,14,15^ Furthermore, whether maladaptive remodeling reflects shared versus phenotype-specific processes across distinct PAH subtypes, such as in SSc-PAH, also remains unclear. Prior molecular profiles of pulmonary hypertensive RV tissue have identified alterations in metabolism, fibrosis, inflammation, and sarcomeric biology, but most have relied on bulk tissue profiling of end-stage specimens obtained via explant or autopsy, lacking cell type-specific resolution and limiting discernment of adaptive versus maladaptive mechanisms from terminal heart failure biology.^16–18^ No prior studies have integrated high- resolution molecular profiling of the living human RV with contemporaneously obtained RV functional measurements to understand how RV cell biology relates to clinically relevant measures of RV adaptation.

In this study, we performed single-nucleus RNA sequencing of human RV endomyocardial biopsies obtained during invasive multi-beat pressure-volume loop assessment in people with idiopathic PAH (IPAH), SSc-PAH, or SSc without PH. By combining RV functional measures with cell type-specific transcriptomic analyses and complementary proteomics, we sought to identify biologic programs associated with RV function across loading conditions. We hypothesized that preserved RV-PA coupling would be characterized by coordinated regulatory and signaling programs involving multiple cardiac cell types.

## Methods

Right ventricular endomyocardial biopsy specimens were obtained from 32 disease subjects with idiopathic PAH (n=12), systemic sclerosis-associated PAH (n=10), and SSc without PH (n=10) during right heart catheterization with contemporaneous RV pressure-volume loop assessment, as previously described.^11,19^ Nonfailing RV tissue from 24 donor hearts suitable but not used for transplantation served as control tissue.^20,21^ Single-nucleus RNA sequencing was performed using the iCell8 platform.^22^ Disease nuclei (IPAH, SSc-PAH, SSc) were assigned subject-level identifiers via genotype-informed demultiplexing with *SoupLadle*;^23^ control nuclei were analyzed at the iCell8 chip level. After quality control filtering, 5,911 nuclei were retained, including 2,273 disease-derived nuclei and 3,638 control nuclei (Supplemental Table 1).

Cells were clustered using *Seurat* and manually annotated based on differential expression of canonical marker genes for each lineage (Supplemental Figure 1). Transcriptional differences within each cell type were modeled using negative binomial mixed models (*nebula* package for R),^24^ with IPAH and SSc-PAH pooled as PAH, and control nuclei serving as the reference group. Because pathway-level analysis can capture coordinated biological changes that may be subtle at the level of individual genes, primary inference focused on pathway enrichment rather than individual-gene differential expression.^25^ Gene set enrichment analysis (GSEA; *fgsea* package for R) was performed using ranked lineage-specific differential expression statistics with Reactome pathways serving as reference groupings for gene sets. Pathway enrichment was summarized using the normalized enrichment score (NES), a size-adjusted enrichment statistic that normalizes for gene-set size and enables comparison of enrichment magnitude across pathways. To evaluate SSc modification of PAH-associated biology, we fit negative binomial models that included PAH status, scleroderma status, and their interaction; sensitivity analyses compared derived contrasts with complementary covariate-adjusted pseudobulk DESeq2 analyses directly comparing IPAH and SSc-PAH.

To link RV transcriptional programs to RV functional measures, lineage-specific pseudobulk expression was regressed on RV pressure-volume measures using age- and sex-adjusted DESeq2 models. Reactome pathway enrichment was performed from ranked Wald statistics.

For a subset of RV biopsies with sufficient tissue available (7 IPAH, 6 SSc-PAH, 5 SSc) and NF controls, bulk proteomic profiling was performed using the Complete360 DISCOVERY platform (Complete Omics Inc., Baltimore, MD), a data-independent acquisition mass spectrometry workflow optimized for solid tissue analysis. Proteomic analyses were performed using a gene-level abundance matrix generated from 3,615 proteotypic (gene-specific) peptides, reducing ambiguity from shared peptide assignments and permitting integration with transcriptomic Reactome gene sets. Protein-level differential abundance analyses compared PAH versus control samples, preserved versus impaired RV-PA coupling, and continuous RV functional measures using appropriate linear models (i.e., *limma)*. Ranked protein-level statistics were tested against Reactome pathways using GSEA (i.e., *fgsea)*. Proteomic enrichments were compared with lineage-resolved RNA gene set enrichments at the Reactome pathway level. Concordant enrichment direction and overlap among pathway leading-edge genes and proteins permitted identification of biologic programs reproducibly represented across molecular layers. False discovery rate correction was applied separately within each predefined inferential family.

A schematic of the overall study design is shown in Figure 1A. All analyses were performed using R version 4.5.2. Detailed tissue processing, demultiplexing, quality control, model specification, and visualization methods are provided in the Supplemental Methods.

**Figure 1.**
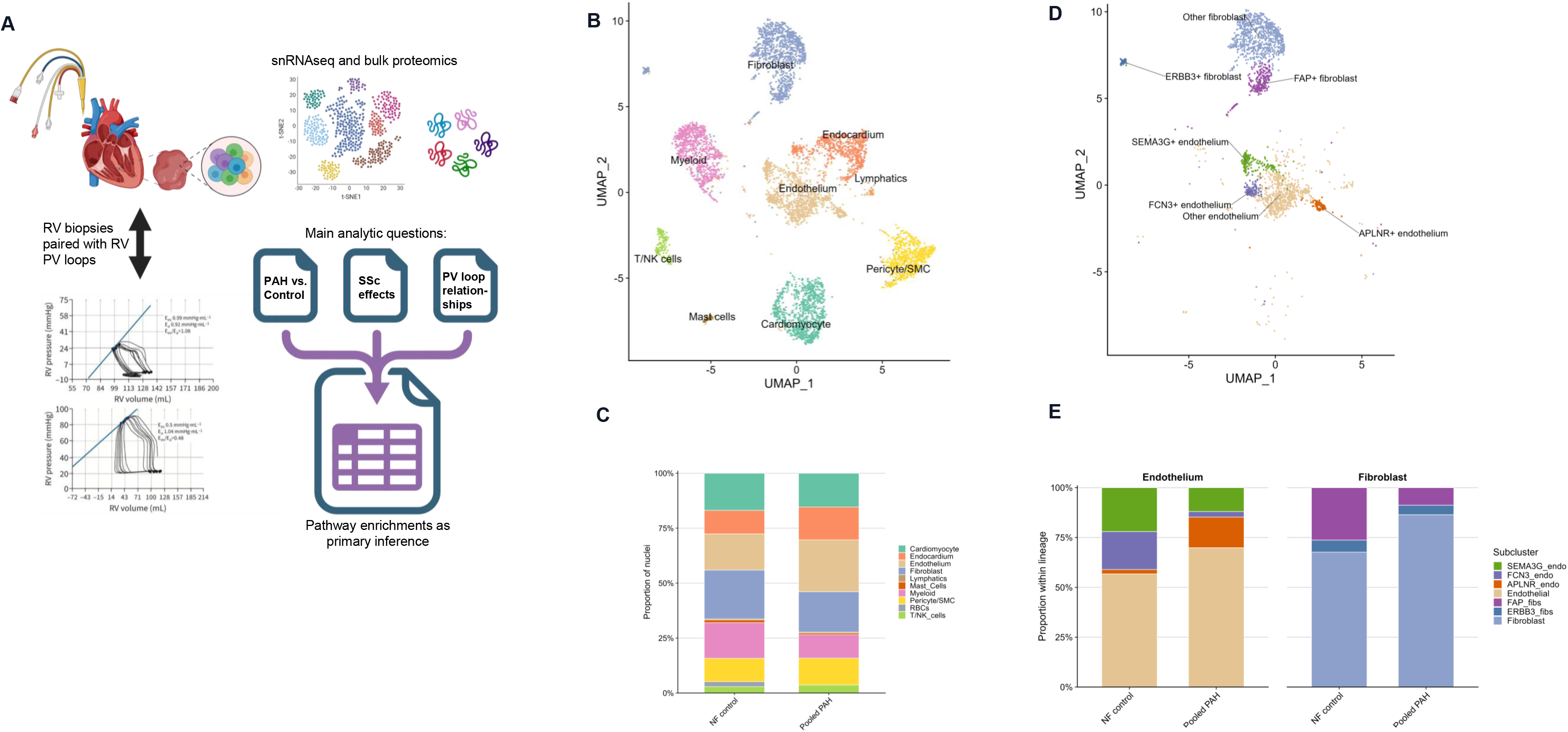
Overall study design and cellular landscape of human right ventricular biopsy tissue in PAH. (A) Schematic overview of study design, integrating right ventricular endomyocardial biopsy tissue obtained contemporaneously with pressure-volume loop measurements, single-nucleus RNA sequencing, bulk proteomics, and pathway-level analyses. (B) UMAP of major right ventricular cell populations. (C) Major-cell-type proportions in nonfailing control and pooled PAH samples. (D) UMAP of endothelial and fibroblast sublineages in nonfailing control and pooled PAH samples. (E) Endothelial and fibroblast subcluster composition in nonfailing control and pooled PAH samples. PAH indicates pulmonary arterial hypertension; PV, pressure-volume; RV, right ventricular; snRNAseq, single-nucleus RNA sequencing; and UMAP, uniform manifold approximation and projection.

## Results

### Cohort demographics and clinical characteristics

As shown in Table 1, individuals with PAH had moderate-to-severe disease, with the majority experiencing NYHA functional class II or III symptoms. All PAH patients were receiving treatment for prevalent disease at the time of biopsy. Individuals with SSc-PAH were older and predominantly female, consistent with the known epidemiology of SSc-PAH.^13^ RV pressure-volume measurements demonstrated reduced RV-pulmonary arterial coupling in SSc-PAH relative to IPAH.^7,10^

**Table 1.** Demographics and Clinical Characteristics of Sequenced Subjects.

| Characteristic | All PAH | IPAH | SSc-PAH | SSc without PH | NF control |
| --- | --- | --- | --- | --- | --- |
| Subjects, n | 22 | 12 | 10 | 10 | 24 |
| Age, y | 59 [56, 68] | 56 [45, 64] | 63 [59, 68] | 52 [39, 69] | 55 [38, 63] |
| Female sex, n (%) | 18 (82%) | 9 (75%) | 9 (90%) | 8 (80%) | 11 (46%) |
| NT-proBNP, pg/mL | 231 [133, 397] (n=19) | 176 [109, 283] (n=10) | 299 [146, 397] (n=9) | 254 [74, 356] (n=9) | — |
| 6-minute walk distance, m | 410 [366, 500] (n=19) | 493 [390, 570] (n=11) | 374 [291, 417] (n=8) | 457 [418, 472] (n=9) | — |
| NYHA/WHO functional class I, n (%) | 4 (18%) | 3 (25%) | 1 (10%) | 1 (10%) | — |
| NYHA/WHO functional class II, n (%) | 13 (59%) | 8 (67%) | 5 (50%) | 7 (70%) | — |
| NYHA/WHO functional class III, n (%) | 5 (23%) | 1 (8%) | 4 (40%) | 2 (20%) | — |
| Right atrial pressure, mm Hg | 6 [5, 8] (n=22) | 5 [4, 6] (n=12) | 6 [5, 8] (n=10) | 4 [4, 6] (n=10) | — |
| Mean pulmonary artery pressure, mm Hg | 36 [29, 40] (n=22) | 38 [30, 42] (n=12) | 36 [26, 40] (n=10) | 16 [14, 17] (n=10) | — |
| Pulmonary capillary wedge pressure, mm Hg | 10 [8, 13] (n=22) | 10 [9, 12] (n=12) | 11 [7, 14] (n=10) | 10 [8, 11] (n=10) | — |
| Thermodilution cardiac output, L/min | 4.5 [3.9, 4.9] (n=22) | 4.5 [4.1, 5.0] (n=12) | 4.3 [3.6, 4.8] (n=10) | 5.9 [4.2, 6.9] (n=10) | — |
| Thermodilution cardiac index, L/min/m<sup>2</sup> | 2.52 [1.99, 2.77] (n=22) | 2.52 [2.00, 2.79] (n=12) | 2.44 [1.82, 2.70] (n=10) | 2.88 [2.40, 3.37] (n=10) | — |
| Pulmonary vascular resistance, WU | 5.92 [3.36, 8.09] (n=22) | 6.02 [4.12, 7.50] (n=12) | 5.77 [3.08, 10.00] (n=10) | 1.30 [0.65, 2.24] (n=10) | — |
| End-systolic elastance (Ees), mm Hg/mL | 0.63 [0.47, 0.90] (n=16) | 0.71 [0.60, 0.94] (n=10) | 0.50 [0.28, 0.76] (n=6) | 0.57 [0.41, 0.85] (n=6) | — |
| Arterial elastance (Ea), mm Hg/mL | 0.76 [0.55, 1.01] (n=17) | 0.77 [0.56, 0.95] (n=10) | 0.66 [0.53, 1.29] (n=7) | 0.29 [0.22, 0.40] (n=6) | — |
| RV-PA coupling (Ees/Ea), ratio | 0.90 [0.58, 1.31] (n=16) | 1.00 [0.81, 1.72] (n=10) | 0.58 [0.45, 1.13] (n=6) | 1.99 [1.45, 2.13] (n=6) | — |
| RV-PA coupling $\geq 1$, n/N (%) | 7/16 (44%) | 5/10 (50%) | 2/6 (33%) | 6/6 (100%) | — |
| RV-PA coupling $< 1$, n/N (%) | 9/16 (56%) | 5/10 (50%) | 4/6 (67%) | 0/6 (0%) | — |
| PDE5 inhibitor, n (%) | 16 (76%) | 11 (92%) | 5 (56%) | 3 (30%)* | — |
| Endothelin receptor antagonist, n (%) | 13 (62%) | 9 (75%) | 4 (44%) | 1 (10%)* | — |
| Prostacyclin pathway agent, n (%) | 1 (5%) | 1 (8%) | 0 (0%) | 0 (0%) | — |

Values are median [IQR] unless otherwise indicated. Clinical and hemodynamic values are shown with the number of subjects with available data. Percentages are calculated among subjects with nonmissing data. Ea indicates arterial elastance; Ees, end-systolic elastance; ERA, endothelin receptor antagonist; IPAH, idiopathic pulmonary arterial hypertension; NF, nonfailing; NT-proBNP, N-terminal pro-B-type natriuretic peptide; NYHA, New York Heart Association; PA, pulmonary arterial; PAH, pulmonary arterial hypertension; PDE5, phosphodiesterase type 5; PH, pulmonary hypertension; RV, right ventricular; SSc, systemic sclerosis; WHO, World Health Organization; and WU, Wood units. \*Indicates prescription of PDE5 inhibitor or ERA for Raynaud phenomenon in SSc. Hemodynamic data are not available for NF controls.

### Clustering and cell-type proportions across conditions

Clustering identified the expected major cardiac cell lineages, including cardiomyocytes, endocardium, endothelium, fibroblasts, pericyte/smooth muscle cells, myeloid cells, and T/NK populations, with clear separation of lineages on low-dimensional embedding (Supplemental Figure 1A and Figure 1B). Relative to controls, PAH samples demonstrated increases in endothelial and endocardial cells, with relative reductions in fibroblasts and myeloid cells (Figure 1C and Supplemental Table 2). Endocardial, endothelial and fibroblast populations together comprised more than half of disease nuclei, therefore we evaluated substructure within these populations.

Coherent sub-lineages formed within fibroblasts and endothelial cells, whereas endocardial nuclei did not resolve into reproducible subclusters (Supplemental Figure 1B and Figure 1D). Subclusters with a canonical marker-defined phenotype were annotated to indicate marker-enriched discrete states, including apelin receptor-enriched (APLNR+), ficolin-3- enriched (FCN3+), and semaphorin-3-G-enriched (SEMA3G+) endothelial populations, and fibroblast-activated-protein-enriched (FAP+) and ERBB3-enriched (ERBB3+) fibroblast populations, while subclusters without a distinguishing marker-defined phenotype retained their parent lineage designation. Within endothelial populations, APLNR+ endothelial cells expanded in disease, while FCN3+ and SEMA3G+ endothelial subclusters showed relative reductions. The FAP+ fibroblast sub-lineage demonstrated a relative reduction in disease (Figure 1E).

### PAH versus control differences

We first applied GSEA to cell type-specific PAH-versus-control differential expression results (Supplemental Figure 2) to define the molecular background upon which RV remodeling occurs. Figure 2A shows transcriptomic GSEA results prioritized by concordant protein-level enrichment. The dominant cross-lineage observation at the RNA level was relative depletion of fundamental cellular maintenance programs in PAH, including RNA processing, translation and ribosome quality control, repair programs, and cell cycle and stress response programs. Pathways related to glucose metabolism and mitochondrial function, such as aerobic respiration and electron transport chain pathways, were also broadly downregulated. Many of these themes were supported by directionally concordant enrichments in the proteomic layer. Most unsupported enrichments had measurable representation in the proteomic dataset, indicating absence of protein-level support was rarely attributable to undetected protein (Supplemental Table 3). Supplemental Figure 3 shows top up- and down-regulated PAH-vs-control transcriptomic enrichments independent of proteomic concordance.

**Figure 2.**
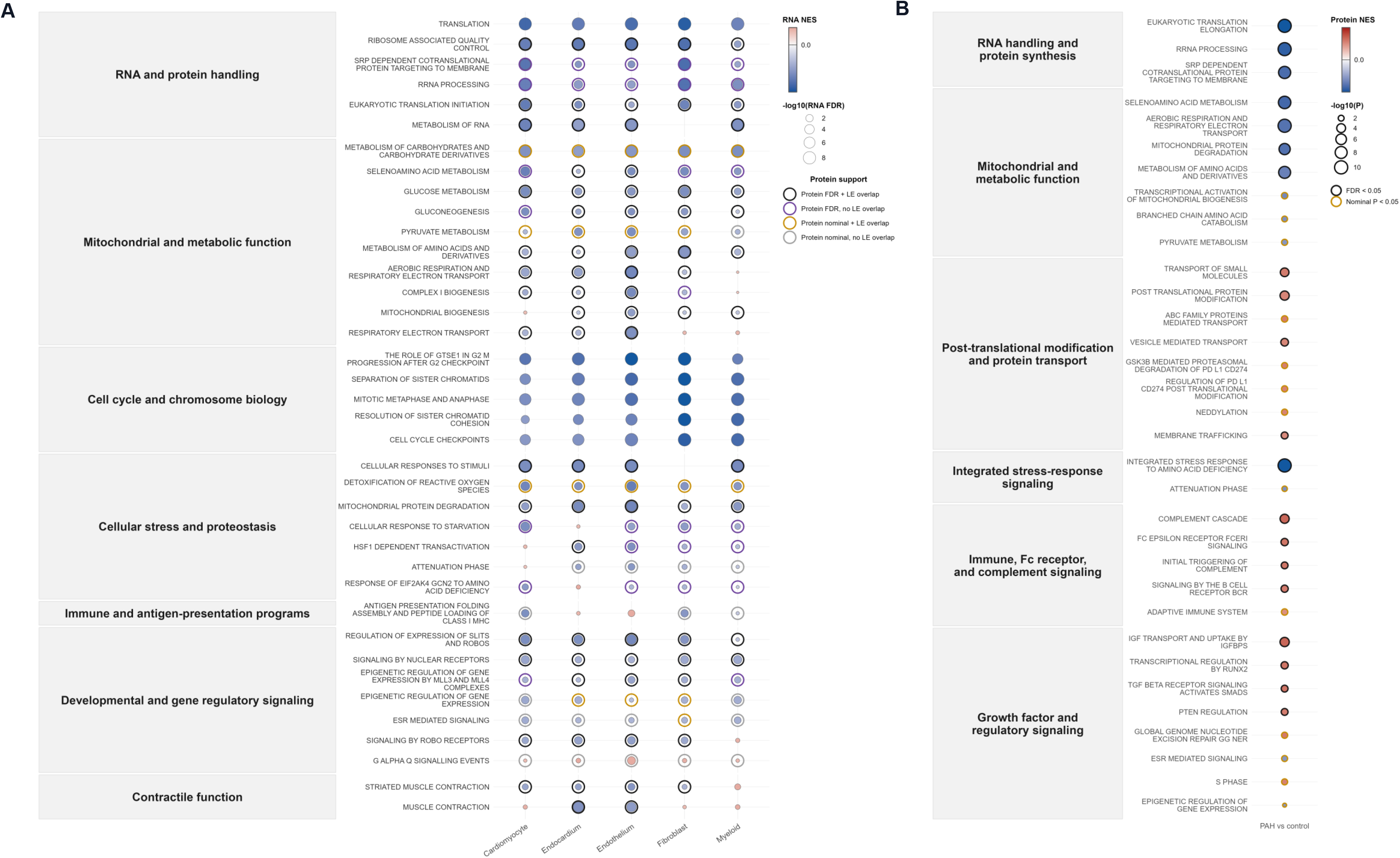
PAH-associated gene and protein set enrichments. (A) Cell type-specific RNA-level Reactome GSEA for PAH versus control. Gene sets were grouped thematically to aid visualization and ordered within themes by overall enrichment pattern. Bubble fill indicates RNA NES, and bubble size indicates −log10(RNA FDR). Outer rings indicate matching bulk RV proteomic GSEA support for the same Reactome pathway: black, protein FDR <0.05 with concordant NES direction and ≥3 overlapping leading-edge genes/proteins; orange, protein nominal P<0.05 with concordant direction and ≥3 leading-edge overlap; purple, protein FDR <0.05 with concordant direction but <3 leading-edge overlap; gray, protein nominal P<0.05 with concordant direction but <3 leading-edge overlap. (B) Bulk RV proteomic Reactome GSEA for PAH versus control. Bubble fill indicates protein NES, bubble size indicates −log10(protein P value), and outer rings indicate protein GSEA support at FDR <0.05 or nominal P<0.05. FDR indicates false discovery rate; GSEA, gene set enrichment analysis; NES, normalized enrichment score; PAH, pulmonary arterial hypertension; and RV, right ventricular.

In protein-only PAH-versus-control enrichments (Figure 2B), RNA handling, protein synthesis, and mitochondrial and metabolic function pathways remained prominent downregulated programs. Upregulated programs included post-translational modification and protein transport programs, Fc receptor and complement signaling, and growth factor and regulatory pathways, including TGF-beta and insulin-like growth factor signaling.

### Decomposition of PAH and scleroderma effects

Given the prevalence of SSc in our cohort, we tested whether SSc modified PAH-associated transcriptional programs by identifying pathways with both significant PAH main effects and significant PAH-by-SSc interactions. Fibroblasts had the highest proportion of SSc- modified PAH-associated pathway calls (58%), followed by cardiomyocytes, endothelial cells, myeloid cells, and endocardial cells. These proportions remained stable in a sensitivity analysis that collapsed redundant leading-edge gene lists (Supplemental Table 4). Across lineages, PAH associations were largely concordant in direction between IPAH and SSc-PAH relative to controls, with interaction effects most often reflecting amplification or attenuation of effect, rather than reversal. Directional discordance was observed in less than two percent of SSc- modified pathways.

The few divergences observed between IPAH and SSc-PAH occurred in biologically plausible pathways, including fibroblast glycosaminoglycan biosynthesis and mitochondrial fatty acid beta-oxidation, cardiomyocyte glycogen metabolism and endothelial carbon and lipid metabolism (all downregulated relative to controls in IPAH, yet upregulated relative to controls in SSc-PAH). Concordant directionality was observed between derived contrasts from interaction models and direct subject-level comparisons, with the exception of cardiomyocyte glycogen metabolism (Supplemental Figure 4B).

### Cell type-specific transcriptional associations with RV function

Having defined the PAH background and its few SSc modifications, we next examined RV biopsy transcriptomic associations with contemporaneously obtained RV pressure-volume measurements (Figure 3). Pathway-level associations were most prominent for the RV-centric measures of RV-PA coupling (Ees/Ea) and contractility (Ees) (Supplemental Figure 5 and Supplemental Table 5), whereas broader hemodynamic indices such as right atrial pressure and thermodilution cardiac output showed comparatively fewer and weaker pathway-level associations despite greater sample completeness (Table 1). RV-PA coupling was associated with transcriptional programs across multiple lineages, with prominent contributions from cardiomyocytes, endothelial cells, endocardial cells, and myeloid cells (Supplemental Figure 5).

**Figure 3.**
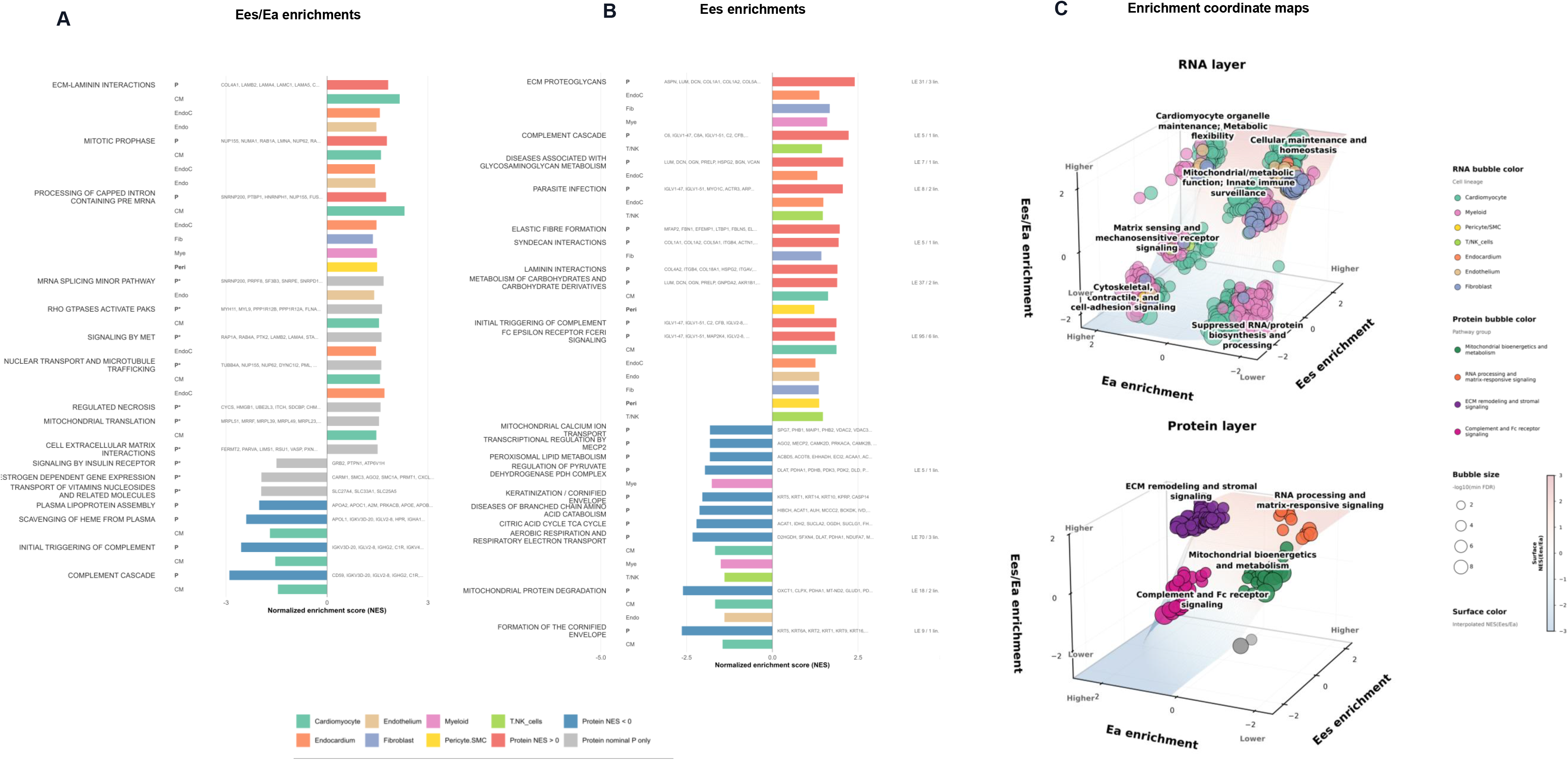
Pathway enrichments associated with RV functional measurements. Bulk proteomic Reactome GSEA associated with continuous Ees/Ea (A) and Ees (B), with corresponding RNA-layer support shown beneath each protein pathway. Protein pathways with nominal protein P<0.05 were collapsed separately by enrichment direction using protein leading-edge Jaccard similarity ≥0.35, with representative pathways manually selected with consideration of protein FDR, protein P value, and absolute protein NES. Colored RNA bars indicate matching lineage-specific RNA enrichments with concordant direction, RNA P<0.05, and ≥3 overlapping non-immunoglobulin leading-edge genes/proteins with minimal proportional overlap. Protein rows are labeled P for protein FDR <0.05 and P* for protein nominal P<0.05. Ees indicates end-systolic elastance; Ea, arterial elastance; FDR, false discovery rate; GSEA, gene set enrichment analysis; NES, normalized enrichment score; PA, pulmonary arterial; and RV, right ventricular. RNA- and protein-level Reactome GSEA results were mapped in a shared physiology-coordinate space defined by normalized enrichment scores (NES) for end-systolic elastance (Ees), arterial elastance (Ea), and RV-pulmonary arterial coupling (Ees/Ea) in (C). Each point represents a lineage-pathway enrichment for the RNA layer or a protein-pathway enrichment for the proteomic layer. Pathways were included if they were significantly associated with at least one of the three traits. Points are plotted by NES for Ees, Ea, and Ees/Ea on the x-, y-, and z-axes, respectively. Dot size reflects the strongest pathway-level significance across the three traits, calculated as −log10 of the minimum FDR. Axis descriptors indicate lower versus higher enrichment along each physiologic dimension. The smoothed surface represents interpolated NES for Ees/Ea and is included as a visual aid for depth and orientation. (Top) RNA-layer enrichments are colored by cell lineage. Recurrent pathway neighborhoods in three-dimensional RNA enrichment space were identified using k-means clustering (k=6) based on NES coordinates for Ees, Ea, and Ees/Ea. Cluster labels summarize dominant biologic themes among representative pathways within each cluster. (Bottom) Protein-layer enrichments were similarly clustered in NES coordinate space and annotated according to dominant biologic themes among member Reactome pathways. Lower-priority or outlier clustered pathways are shown in gray. For both layers, post hoc analyses were used to assess the biologic distinctiveness of pathway clusters. Leading-edge gene composition differed significantly across RNA-layer clusters by PERMANOVA on Jaccard distances, including after permutation stratified by lineage (P=0.001), and across protein-layer clusters (P=0.001). Preserved-coupling regions with lower versus higher Ees/Ea also differed in leading-edge composition at both the RNA and protein layers (P=0.001 for each). Pairwise leading-edge redundancy was low overall, with median within-cluster Jaccard similarity of 0.00 in both layers, supporting that cluster structure was not driven solely by duplicate Reactome gene sets. Theme-permutation, preserved-region theme tests, and leading-edge overrepresentation analyses are provided in the accompanying source workbook.. Ees indicates end-systolic elastance; Ea, arterial elastance; FDR, false discovery rate; NES, normalized enrichment score; and RV, right ventricular.

With enrichment against continuous RV-PA coupling, (Figure 3A and Supplemental Figure 6), higher Ees/Ea was associated with upregulated collagen and laminin interactions, extracellular matrix (ECM) interactions, RNA processing programs, and Rho GTPase programs at the protein level, with supporting transcriptomic enrichments observed primarily in cardiomyocyte, endocardial, and endothelial transcriptomes. Lower Ees/Ea was associated with prominent protein-level enrichment of immune pathways, including immunoglobulin-containing, Fc receptor activation, and complement pathways. Protein-level enrichments involving immunoglobulins did not demonstrate overlapping RNA-level enrichments in individual cell types, though complement-triggering and complement cascade pathways exhibited RNA-layer support from cardiomyocytes.

Higher Ees was associated with upregulated ECM proteoglycans, collagen biosynthesis and assembly programs, upregulated syndecan and integrin interactions, and upregulated glycosaminoglycan metabolism, with these protein-level enrichments matched mostly to fibroblast, endocardial, and cardiomyocyte transcriptomes (Figure 3B and Supplemental Figure 7). Lower Ees was associated with enrichments in mitochondrial function programs, including cristae formation, aerobic respiration and electron transport chain, TCA cycle and pyruvate dehydrogenase complex, and formation of ATP. A plurality of protein-level enrichments against lower contractility matched transcriptomic enrichments in cardiomyocytes.

Because Ees and Ees/Ea analyses implicated overlapping pathways with differing enrichment magnitudes and directions, we first plotted NES(Ees) against NES(Ees/Ea) (Supplemental Figure 8). Mitochondrial pathways, for example, were positively associated with Ees/Ea but negatively associated with Ees. Because Ees/Ea reflects contractile performance relative to arterial load, we reasoned that such discordance could reflect differences in afterload. We therefore incorporated pathway enrichment for arterial elastance, NES(Ea), as a third coordinate to assess how molecular programs varied jointly with afterload, contractile response, and RV-PA coupling. This representation resolved clusters of pathways occupying different regions of physiologic coordinate space (Figure 3C).

At the transcriptomic level, cardiomyocyte-predominant mitochondrial and metabolic pathways occupied lower-afterload, lower-contractility, higher-coupling space, whereas lower- afterload, lower-contractility, lower-coupling space was occupied by downregulated RNA and protein biosynthesis and processing programs. Different pathways occupied higher-load, lower- contractility, lower-coupling space (downregulated contractile and cell adhesion signaling programs) versus higher-load, higher-contractility, lower-coupling space (matrix sensing and mechanosensitive receptor signaling programs). Cardiomyocyte-predominant programs involving organelle maintenance and metabolic adaptation, including pyruvate metabolism and fatty acid oxidation pathways, localized to the region associated with higher load, higher contractility, and higher RV-PA coupling.

Plotting proteomic NES coordinates in the same manner demonstrated similar organization of pathway enrichments into clusters. Mitochondrial and metabolic protein-layer programs localized to a similar region as their analogous RNA-layer programs, demonstrating associations with lower load, lower contractility, and higher coupling. Immune and humoral pathways, including Fc gamma receptor signaling and complement activation programs, were associated with higher load, higher contractility, but lower coupling. A distinct protein cluster comprising RNA processing and matrix-responsive signaling localized to lower-load, higher- contractility, higher-coupling space. Upregulated collagens, laminins, integrins, and ECM pathways were associated with higher load, higher contractility, and higher coupling.

To place pathway associations with RV-PA coupling into a clinically anchored framework, we next compared pathway enrichments between individuals with coupled versus uncoupled RV-PA units (Ees/Ea ≥1 vs <1). The dichotomized Ees/Ea analysis resolved the gradients observed in continuous models into two multicellular RV states. As shown in Figure 4A, at the RNA level, preserved coupling (with positive NES) was characterized by non-myocyte enrichment of ECM organization pathways and RNA processing and translational machinery programs, with signal localizing primarily to endocardial cells, endothelial cells, and fibroblasts. An analogous proteomic analysis dichotomized by RV-PA coupling showed enrichments for integrin-cell surface interactions, mitochondrial translation, and laminin and ECM proteoglycan programs in coupled RVs, with RNA-layer endocardial support for several matrix-associated programs, and endothelial support for integrin-cell surface interactions (Figure 4B). Lineage-pathway NES remained stable in sensitivity analyses in which coupling associations were adjusted for PAH subtype (IPAH vs. SSc-PAH), and in which the coupling threshold was shifted to Ees/Ea ≥0.8 vs <0.8 (Supplemental Figure 9).

**Figure 4.**
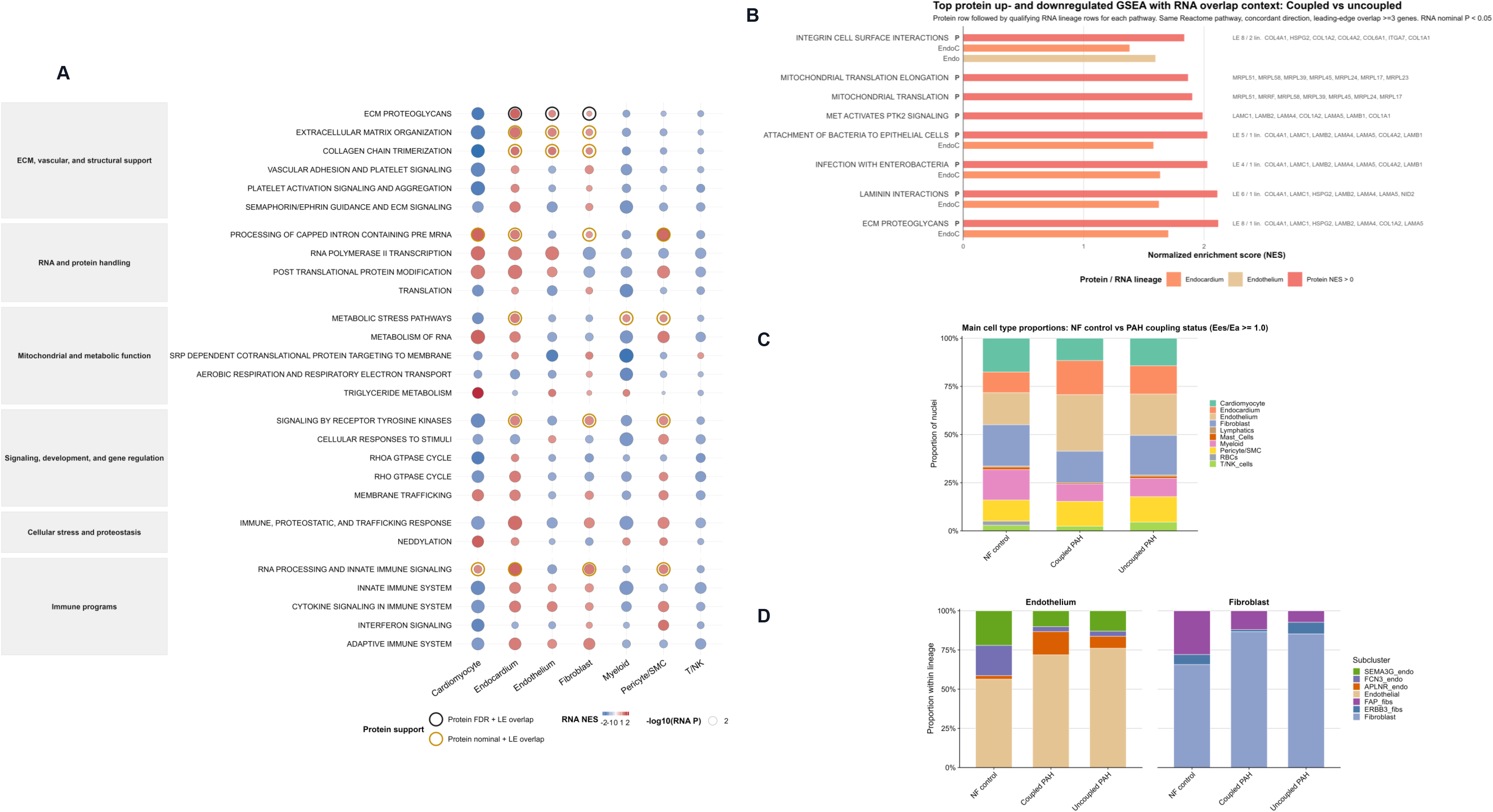
Cellular and pathway-level features of preserved versus impaired RV-PA coupling. RV-PA coupling status was dichotomized as preserved coupling (Ees/Ea ≥1) or impaired coupling (Ees/Ea <1). (A) RNA-level Reactome GSEA comparing preserved versus impaired coupling, prioritized from the original RNA-first/protein-support display and collapsed by leading-edge gene similarity. Displayed pathways were clustered using Jaccard similarity of RNA leading-edge genes with a threshold of 0.35; one representative pathway was retained per cluster, prioritizing protein support, number of supported lineages, RNA P value, and RNA NES magnitude. Bubble fill indicates RNA NES, bubble size indicates −log10(RNA P value), and outer rings indicate concordant bulk proteomic support using the same criteria as in Figure 2. (B) Bulk proteomic Reactome GSEA comparing preserved versus impaired coupling, with corresponding RNA-layer support by cell type. (C) Major-cell-type proportions across nonfailing control, coupled PAH, and uncoupled PAH RVs. (D) Endothelial and fibroblast subcluster composition across coupling groups. Ees indicates end-systolic elastance; Ea, arterial elastance; FDR, false discovery rate; GSEA, gene set enrichment analysis; NES, normalized enrichment score; PAH, pulmonary arterial hypertension; and RV, right ventricular.

At the level of main cell types, coupled RVs showed relatively increased endocardial and endothelial populations compared to controls, though no differences were statistically significant when coupled vs. uncoupled RVs were directly compared (Figure 4C and Supplemental Table 2B). At the subpopulation level, coupling was associated with expansion of the *APLNR*+ endothelial subcluster (Figure 4D).

### Predicted cell-cell signaling patterns

Given the multicellular nature of the programs we identified as associated with RV-PA coupling, we performed exploratory cell-cell signaling analyses in PAH RVs stratified by coupling status, with NF control signaling also analyzed as a reference (Figure 5 and Supplemental Figures 10 and 11). Inferred interactions in coupled RVs were dominated by ECM-associated signaling, with fibroblasts prominent as sending populations (Figure 5A). Several interactions present in nonfailing RVs demonstrated greater inferred signaling probability in coupled PAH RVs, including PECAM and ESAM homotypic adhesion interactions among endothelial and endocardial cells, and fibroblast-derived laminin-integrin signaling (Supplemental Figure 10B). PDGFB-PDGFRB signaling toward smooth muscle cells, absent in control RVs in this analysis, emerged in PAH and demonstrated greater inferred probability in coupled than uncoupled RVs (Figure 5B). Broadly, coupled RVs demonstrated predicted laminin-integrin interactions across fibroblast, cardiomyocyte, endothelial, and mural-cell compartments, including fibroblast-derived laminins and collagens paired with integrin receptors in endothelial, smooth muscle, and cardiomyocyte populations.

**Figure 5.**
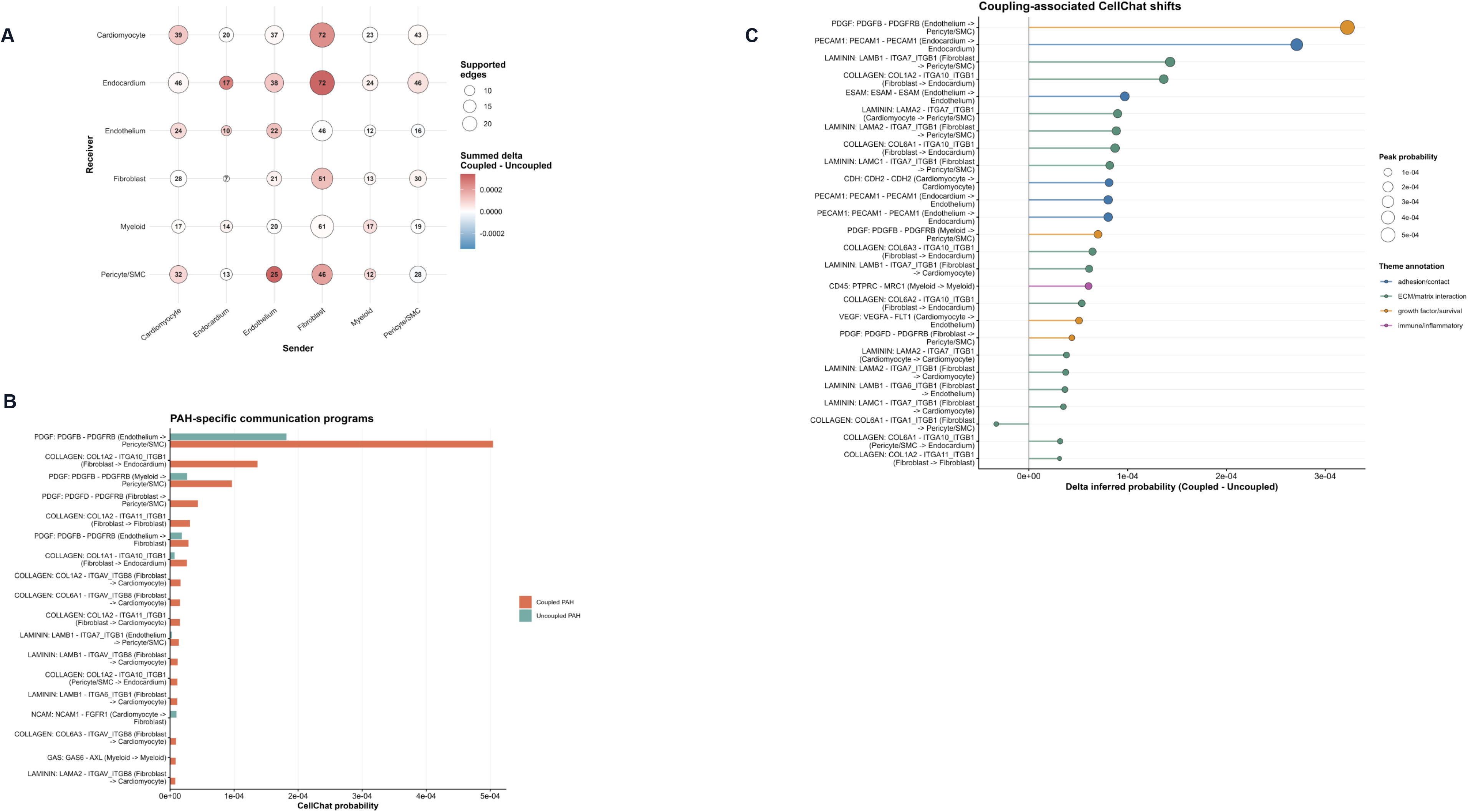
Predicted cell-cell communication patterns in coupled versus uncoupled PAH RVs. Exploratory CellChat analyses were performed in PAH RVs stratified by preserved coupling (Ees/Ea ≥1) versus impaired coupling (Ees/Ea <1), with nonfailing control RVs analyzed as a reference. (A) PAH-specific communication programs are shown according to CellChat pathway probability in coupled and uncoupled RVs. (B) Coupling-associated CellChat shifts are shown for selected ligand-receptor pairs, with point size indicating peak inferred probability and color indicating signaling category. (C) Sender-receiver matrix summarizing directional changes in communication strength between major cell compartments in coupled versus uncoupled RVs. Ees indicates end-systolic elastance; Ea, arterial elastance; PAH, pulmonary arterial hypertension; RV, right ventricular; and SMC, smooth muscle cell.

To provide independent spatial context for these predicted interactions, we examined segmented (fibroblast, vascular, or cardiomyocyte) RV regions of interest in a human heart-failure RV GeoMx dataset (GSE271676). At the individual gene level, ECM support transcripts recurrently implicated in our analyses localized to fibroblast-associated segments. *PECAM1* and *ESAM* colocalized with the endothelial marker *VWF* in vessel-adjacent segments (Supplemental Figure 12A). When summarized as composite expression scores for predefined ECM-support and endothelial adhesion gene sets, the ECM-support genes implicated recurrently in our results localized preferentially to fibroblast-associated segments, whereas adhesion genes localized to vessel-enriched regions (Supplemental Figure 12B).

## Discussion

To our knowledge, this is the first study to molecularly profile the living human PAH RV at single-cell resolution in direct association with contemporaneously obtained multi-beat pressure-volume measurements of RV function. By integrating snRNA-seq and complementary proteomics with measures of RV afterload (Ea), contractility (Ees), and RV-PA coupling (Ees/Ea), we identified multicellular programs associated with different RV physiologic states. First, we found that PAH was characterized by broad depletion of fundamental biosynthetic, proteostatic, mitochondrial, and repair programs across cardiac cell types. Second, we found preserved RV-PA coupling in PAH was associated with coordinated non-myocyte ECM and vascular-stromal programs in addition to cardiomyocyte metabolic programs. Third, we found the biology associated with preserved coupling differed according to loading conditions and contractile response, with mitochondrial and metabolic competency predominating with relatively lower afterload, and ECM remodeling and vascular support signaling predominating with higher afterload and higher contractility. Together, these findings suggest that RV adaptation is not a single molecular state but a coordinated, multicellular response that varies along with the physiologic demands placed on the RV.

The dominant PAH-associated transcriptional signal was relative depletion of biosynthetic, proteostatic, trafficking, and repair programs across multiple cell types, with superimposed cell type-specific remodeling in structural cells and cardiomyocytes, and a more selective immune-surveillance and interferon phenotype in immune populations. Themes we later found associated with RV-PA coupling, including RNA and protein handling, metabolic flexibility, and mitochondrial function, were downregulated in the initial PAH-to-control comparison, suggesting RV adaptation to load may involve preservation or reacquisition of biologic programs broadly downregulated in PAH. Because the RV tissues in our study were obtained from living patients instead of end-stage specimens, these PAH-versus-control signatures likely capture chronic RV remodeling, rather than terminal heart failure biology. Importantly, recurrent PAH-associated themes had pathway-for-pathway support from protein set enrichments, demonstrating reproducible biology across molecular layers.

The presence of SSc modified the PAH-associated molecular background but did not appear to create an entirely distinct RV response. Although many PAH-associated pathways demonstrated significant PAH-by-SSc interactions, these effects overwhelmingly reflected amplification or attenuation of enrichments rather than directional reversal. A small set of pathways demonstrated true subtype divergence, involving biology plausibly relevant to SSc, including fibroblast extracellular matrix remodeling and fatty acid oxidation, endothelial carbon and lipid metabolism, cardiomyocyte metabolic substrate handling, and selected immune programs. Together, these findings suggest that SSc subtly modifies the structural, metabolic, and immune contextual background for RV remodeling in PAH, rather than fundamentally altering remodeling programs.

The most novel aspect of this study is the integration of RV multi-omic profiles from living humans with contemporaneous pressure-volume loop measures of RV function. RV transcriptional enrichments across multiple cell types were linked closely with RV-PA coupling, supporting the concept that maintenance of coupling requires a coordinated multicellular tissue state rather than simply a cardiomyocyte-intrinsic response to pressure overload. Preserved coupling was characterized by upregulated ECM organization, laminin-integrin signaling, receptor tyrosine kinase signaling (with a strong *PDGFB-PDGFRB* predicted binding signal), and enriched RNA processing and translational capacity across non-myocyte populations. Consistent with these pathway-level patterns, cell type composition analyses showed endocardial and endothelial populations were expanded in the coupled PAH RV, as was observed in PAH overall. There was a trend toward relative depletion of endothelial and endocardial populations in uncoupled RVs, potentially consistent with lost or failed vascular-stromal support.

Our cell-cell signaling predictions further implicate ECM structural support and vascular-stromal signaling in successful coupling. Coupling-associated shifts were concentrated in fibroblast and endothelial senders and endothelial, endocardial and cardiomyocyte receivers. Laminin-binding α7β1 integrins, established mediators of basement membrane adhesion and mechanotransduction,^26,27^ recurred among predicted receivers. The highest-probability coupling-associated interaction was endothelial-to-pericyte/SMC PDGFB–PDGFRB signaling, a recognized regulator of pericyte recruitment and vascular stabilization.^28,29^ Endothelial and endocardial PECAM and ESAM, predicted to bind homotypically, are established junctional adhesion molecules involved in vascular barrier organization and adhesion interactions.^30,31^ In an independent human RV spatial transcriptomic dataset, ECM-support genes implicated by these analyses localized preferentially to fibroblast-associated regions, whereas endothelial adhesion genes localized to vascular regions, demonstrating spatial compatibility of the predicted interactions.

The ECM-associated signal in our results should not necessarily be interpreted as fibrosis alone. The prominence of laminin-integrin, syndecans, and receptor tyrosine kinases alongside collagen and ECM proteoglycan programs in coupled RVs implicates the matrix as both a structural and signaling compartment. Longstanding work suggests integrins are key mechanotransduction receptors linking the extracellular matrix to cardiomyocyte structure and signaling.^26,27^ Classic and contemporary studies have emphasized that integrin-associated complexes regulate force transmission, sarcolemmal stability, costamere organization, and cellular responses to mechanical load.^32–34^ Further, evolving literature shows that cardiac fibroblasts are not merely passive matrix-producing cells, but active regulators of tissue architecture, immune signaling, and paracrine communication.^35^ Recent single-cell and spatial studies across various models of myocardial injury and heart failure have identified fibroblast-immune interactions and fibroblast-endothelial interactions as important components of mechanosensitive ECM programs.^36–39^

Our endothelial sub-lineage signaling was exploratory, though coupling-associated interactions preferentially involved the APLNR+ (apelin-enriched) endothelial subpopulation found to be expanded in PAH, including predicted endothelial PDGFB signaling toward pericyte/SMC populations (Supplemental Figure 10). Prior studies have shown that APLNR is induced in myocardium in response to ventricular loading, and that APLNR+ endothelial cells may promote angiogenesis and myocardial metabolic flexibility through regulation of tissue fatty acid uptake.^40,41^ These signaling predictions support the possibility that maintenance of coupling involves specialized endothelial and stromal support functions that match bioenergetic supply to increased demands. However, because sub-lineage analyses were based on relatively small numbers of nuclei, these findings should be considered hypothesis-generating.

A key observation from our analyses was that interrogation of RV-PA coupling alone did not define a single adaptive biologic state. Preserved coupling can arise from relatively modest contractility matched to lower afterload, or from augmented contractility sufficient to match substantially greater afterload. Mapping pathway associations jointly across Ea, Ees, and Ees/Ea revealed biologic differences underlying these physiologic differences. With relatively lower afterload and contractile demand, preserved coupling was characterized predominantly by cardiomyocyte mitochondrial bioenergetic and metabolic programs, suggesting maintenance of energetic competency in an RV that can match its load without marked augmentation of contractility. In contrast, preserved coupling at higher afterload and Ees was associated with ECM remodeling and stromal signaling programs alongside metabolic flexibility programs. These findings suggest RV adaptation to PAH requires cellular competencies that are engaged depending on load, with increasing afterload requiring additional matrix and mechanosensory support to maintain force transmission across the ventricular wall.

This demonstrated importance of matrix-mediated signaling to RV adaptation may have therapeutic implications, as experimental studies support the concept that ECM cues can be manipulated to improve myocardial function. In a rat model of chronic RV pressure overload induced by pulmonary artery banding, injection of decellularized myocardial matrix hydrogels improved RV systolic function, reduced adverse remodeling and interstitial fibrosis, and enhanced neovascularization.^42^ Though never studied in the human RV, the safety and feasibility of transendocardial injection of matrix hydrogels has been demonstrated in post-myocardial infarction patients with left ventricular dysfunction.^43^ In more targeted experiments, augmentation of laminin–α7β1-integrin signaling improved myocardial tissue integrity in experimental muscular dystrophy,^44,45^ while laminin-221, an α7β1 ligand, enhanced mechanical and mitochondrial function of human induced-pluripotent-stem-cell-derived cardiomyocytes and improved engineered cardiac tissue performance in experimental ischemic cardiomyopathy.^46^ Considered alongside the ECM signaling and laminin-integrin programs identified in our results, these observations raise the possibility that selective reinforcement of matrix-receptor signaling could represent a RV-directed therapeutic strategy in PAH.

Several limitations are acknowledged. First, this study is cross-sectional and therefore cannot establish temporal or causal relationships between transcriptional programs and RV function. Second, although the use of living human RV biopsies represents a major strength, the limited tissue quantity obtainable via this method constrains cell yield and limits assessment of cell subpopulations. A detailed assessment of immune cell types is particularly limited here. Third, due to ethical and practical limitations, nonfailing control tissue was obtained from donor hearts not used for transplantation rather than from biopsies of healthy living subjects. Fourth, proteomic analyses were performed on bulk tissue and therefore lacked cell type-specific resolution. Finally, while we attempt to provide useful spatial contextualization of the programs identified in our data, the external dataset we analyzed for this purpose lacks invasive phenotyping, and therefore we cannot generalize inferences regarding RV function.

In summary, our integrated single-nucleus transcriptomic, proteomic, and physiologic analyses identify RV adaptive responses to PAH as coordinated multicellular processes, rather than a cardiomyocyte-restricted response to pressure overload. Preserved RV-PA coupling was associated with distinct molecular programs according to load and contractile state, ranging from mitochondrial and metabolic competency under lower load, to ECM remodeling and stromal communication under greater load and contractile demand. Together, these findings provide a human molecular framework in which RV adaptation reflects energetic, structural, and vascular-stromal support programs, and identifies candidate biology for future RV-directed therapeutic investigation in PAH.

